# Cortical gray matter density at age five associated with preceding early longitudinal language profiles: A Voxel-based morphometry analysis of the FinnBrain Birth Cohort Study

**DOI:** 10.64898/2026.03.27.714719

**Authors:** Essi Saloranta, Jetro J Tuulari, Elmo P Pulli, Hilyatushalihah K Audah, Aaron Barron, Ashmeet Jolly, Aylin Rosberg, Isabella L.C. Mariani Wigley, Kiia Kurila, Akie Yada, Aura Yli-Savola, Satu Savo, Eeva Eskola, Michelle Fernandes, Riikka Korja, Harri Merisaari, Ekaterina Saukko, Venla Kumpulainen, Anni Copeland, Eero Silver, Hasse Karlsson, Linnea Karlsson, Elina Mainela-Arnold

## Abstract

Previous studies exploring the connection between early language development and brain anatomy have shown that cortical areas relating to individual differences in language skills are diverse and vary depending on the age of child. However, due to lack of large longitudinal samples, current literature is limited in answering the extent to which individual differences in language development prior to school age are reflected in areas of the cortex. To fill this gap, we compared gray matter density between participants that belonged to different longitudinally defined language profiles from 14 months to five years of age in a large population-based sample.

Participants were 166 children from the FinnBrain Birth Cohort Study who had longitudinal language data from 14 months to five years of age and magnetic resonance imaging data at five years of age. Three groups of language development were used as per our prior study: persistent low, stable average, and stable high. Voxel-based morphometry metrics were calculated using SPM12 and the three language profile groups were compared to one another. Covariates included sex and age at brain scan. The statistics were thresholded at p < 0.01 and *false discovery rate* corrected at the cluster level.

Of the three longitudinal language profiles, the stable high group had higher gray matter density than the persistent low group in the right superior frontal gyrus. No differences were found between the stable average and stable high groups, nor persistent low and stable average groups.

The identified superior frontal cortical area belongs to executive functions neural network. This finding adds to the cumulating evidence that individual differences in language development are reflected in growth of gray matter supporting general processing ability rather than specialized language regions. The results suggest that cognitive development and early language development are linked through shared principles of neural growth, identifiable already at age five.

**Key points:**

1. An association between early language development from 14 months to five years of age and gray matter density differences of the right superior frontal gyrus was found at the age of five years. Children following the strongest language trajectory were more likely to exhibit higher gray matter density of the right superior frontal gyrus than children following the weakest trajectory.
2. As the superior frontal gyrus is part of executive functions network, we propose that individual differences in early language development are more defined by general learning mechanisms supported by those networks, rather than language specific pathways.

## 1 Introduction

Brain structures and language undergo largest growth spurts simultaneously in early childhood (Johnson, 2001; Koyama, 2025). The fact that these phenomena co-occur raises questions about their interrelatedness, such as which affects the other and how large the effects are. The dual route hypothesis proposes that dorsal and ventral language routes consisting of areas of gray and white matter work in unison to support language processing (Friederici & Gierhan, 2013). Many existing studies have explored the relationship between language and brain growth (e.g. Girbau-Massana et al., 2014; Pigdon et al., 2019), showing that connections between various cortical and subcortical areas are associated with different aspects of language, especially in adulthood (Friederici, 2011; Richardson & Price, 2009). Fewer studies have been conducted in young children comparing at-risk language development with typical and strong, leaving open which key brain areas might prove essential in developmental outcomes (Herbert, 2004; Soriano-Mas et al., 2009, for more studies see Supplementary material 1). To better answer this question, more studies are needed with participants under the age of five, which is the youngest age at which existing voxel-based morphometry analyses on language development have been successfully conducted (see Supplementary material 1). The sparsity of studies in preschool children may in part be due to the unique challenges in conducting magnetic resonance imaging (MRI) in young children. The current study aims to fill the gap in the literature by associating gray matter (GM) density at age five with early emergence of individual differences in language development, including an at-risk category, collected at FinnBrain Birth Cohort Study in Southwest Finland in a relatively large sample of 166 participants.

### 1.1 Connection between language development and brain development in early childhood

Language develops rapidly from the cries of a newborn into first words of a one-year-old which in turn develop into word combinations of a two-year-old (Määttä et al., 2012). These first combinations gain length and complexity so that by age five the foundations of language that resemble adult-like skill have been set (Lonigan & Milburn, 2017). Majority of children develop rapidly while some experience difficulties that may be transient or persistent in nature (Rescorla, 2011). A great deal of variance is present in early language skills, for example in the size of expressive vocabulary and production on grammatical structure (Huttenlocher et al., 2010). Language difficulties in early childhood are diverse, some limited to small vocabulary (Henrichs et al., 2013), while others cross the whole of linguistic capacity, including difficulties in language comprehension in addition to limited expressive skills (Tomblin et al., 1996). Longitudinal studies on first language acquisition have separated typical development from delayed and precocious development with either using cut-off scores on standardized language tests (Henrichs et al., 2011) or dividing large cohorts into subgroups of language development with data-driven methods (Ukoumunne et al., 2012). A group of persistent language difficulties is almost always present, indifferent from the categorization method, which underlines the importance of identifying factors that associate with this type of language development. This study utilized latent language profiles created with a data-driven method in the FinnBrain Birth Cohort (Saloranta et al., 2025). The group of persisting language difficulties observed in the FinnBrain data resembled two risk categories of language development in a longitudinal continuum: small expressive vocabulary and few word combinations of late talking toddlers at age 30 months continuing to language difficulties at age 5 resembling developmental language disorder (DLD). Late talkers are toddlers aged two to three who lack in vocabulary and word combinations (Rescorla, 2011) whereas DLD is a condition where language difficulties hinder communication in everyday settings in children age 5 and older (Bishop et al., 2017).

The first five years of life when language undergoes a vast growth spurt are also a highly dynamic period for human brain development. Although the expansion of brain volume driven by neurogenesis is complete by mid-gestation, other neurodevelopmental processes peak in the first few postnatal years, including synaptogenesis and myelination, which are responsible for a profound remodelling of cerebro-cortical and white matter microstructure (Budday et al., 2015; Zeiss, 2021). Correspondingly, the rate of global gray matter growth is at lifetime peak between the first and third postnatal years. By five years of age, the cortex is close to its maximum volume (Bethlehem et al., 2022). This period also witnesses a substantial increase in integrity of white matter fibre orientation, driven primarily by the high rate of myelination (Lebel & Deoni, 2018). Thus, by five years of age, the brain has undergone considerable structural remodelling. In line with this, there is a functional reorganization at this age, as the brain begins to exhibit increasing functional lateralization in language-associated regions during language tasks (Olulade et al., 2020).

### 1.2 Voxel-based morphometry analyses in language development research

Evidence from the past decades indicates that two anatomically and functionally distinct pathways, the dorsal and ventral pathways, support language processing in adults and children (see, Friederici & Gierhan, 2013). The dorsal language pathways include gray matter areas of the pars opercularis, the inferior frontal gyrus, the transverse temporal gyrus, the superior temporal gyrus, planum temporale, and the supramarginal gyrus, with white matter tracks connecting temporal cortex to frontal cortex via the arcuate fasciculus and superior longitudinal fasciculus. In comparison, the ventral language pathways include gray matter areas of the inferior temporal gyrus, the middle temporal gyrus, the temporal pole, the pars triangularis and the pars orbitalis of the inferior frontal gyrus, with white matter tracks connecting the middle temporal lobe and the ventrolateral prefrontal cortex via the extreme capsule. Although recent studies suggest structural differences in dorsal and ventral language pathways in young adults with DLD when compared to peers (e.g. Lee et al., 2020), all in all, studies comparing children with DLD to peers have also yielded numerous other differences. These involve differences in overall gray matter volume (Girbau-Massana et al., 2014), overall white matter volume (Herbert et al., 2003), volumes of subcortical (Lee et al., 2013) and several regional cortical areas (Badcock et al., 2012; Soriano-Mas et al., 2009), as well as overall white matter integrity (Verly et al., 2019). However, to our knowledge, past research has not attempted to examine the extent to which the longitudinally emerging early language risk associates with structural differences in the developing brain.

Gray matter density represents the probability that a given voxel contains gray matter rather than white matter (Ashburner & Friston, 2000). In comparison to region-of-interest based analysis, voxel-vise analyses on density map statistically significant clusters anywhere in the brain. This is particularly useful in the study of language development, where potential associations have been reported in many brain regions, such as the fronto–parietal cortex (Badcock et al., 2012), the occipital cortex (Girbau-Massana et al., 2014), the cerebellum (Pigdon et al., 2019) and the subcortical system (Soriano-Mas et al., 2009). The direction of the association changes over the course of development as the generation of new connections before the volume peak gives way to synaptic pruning that better describes refined development in later years. Indeed, in previous studies focusing on older children and teens after the GM growth peak of early childhood, higher volumes have been associated with weaker level of skill (cf. Supplementary Material 1). Studies that focus on early stages of language development are currently lacking.

The current study aims to fill the gap in the literature by providing information on the emergence of individual differences and risk in language development on gray matter density at age five years. Prior longitudinal studies on language and brain development have mostly been conducted in older children, giving little information on the relationship between gray matter structure and individual differences in language development in its early stages. As five-year-olds are old enough to co-operate during imaging, but young enough to not have enrolled into the school system in Finland, the sample we used has a unique potential in answering the question of the extent to which individual differences in early language development can be linked to gray matter structures. This study aims to answer the following research questions: 1) Are there differences in cortical gray matter density at age five between children belonging to the three longitudinal language profiles observed in the FinnBrain cohort? and 2) If so, which cortical areas differ between the developmental language groups? As previous studies have mixed findings about which cortical areas link to individual differences in early language development, and the direction of these associations, we set to explore the connections in our data on whole-brain level voxel-vise morphometry. We hypothesized that children following different early language trajectories would differ in the relative of volume of the language areas of the left hemisphere as opposed to relative volumes of other areas such as fronto-temporal, occipital, subcortical and cerebellar areas.

## 2 Materials and Methods

Data for the study were derived FinnBrain Birth Cohort Study, a longitudinal research effort to detangle how biological and environmental factors affect developmental outcomes (Karlsson et al., 2018). The participating families were recruited on gestational week 12, with 3939 families recruited overall. The original sample is representative of the Finnish population (Karlsson et al., 2018). Follow-up of the cohort consisted of questionnaires sent to everyone, with smaller subsamples invited to laboratory measurements concerning, for instance, language measurements and brain imaging.

### 2.1 Participants

The sample for this study consisted of children who were included in the longitudinal language profile data (n = 1281) and who participated in the five-year-old MRI data collection (n = 173). The inclusion criteria applied to the longitudinal language modeling were the following: 1) participated in at least one language measurement at 14 months, 30 months or five years of age and 2) language exposure to Finnish language at least 80 percent of waking time based on parental report. The exclusion criteria for the five-year-old neuroimaging visit were 1) premature birth (< 35 gwks; < 32 gwks for those participating in a nested substudy examining the effects of maternal prenatal synthetic glucocorticoid treatment), 2) developmental sensory deficits or congenital anomalies (e.g., blindness, congenital heart disease), 3) severe chronic pediatric and child neurological conditions (e.g., epilepsy), 4) current ongoing medical examinations or clinical follow-ups (i.e., referrals from primary to specialized medical care such as hearing level evaluations). Clinically identified DLD was neither an inclusion nor an exclusion criterion. 5) regular daily medications (e.g., oral medications, inhalants and topical creams. An exception to this was desmopressin (®Minirin)), 6) history of head trauma and 7) ear tubes and routine MRI contraindications. Only those who had both longitudinal language data and a successful MRI scan were included in this study. Overlap between the longitudinal language and the cross-sectional MRI datasets resulted in a final sample size of n = 166.

### 2.2 Procedures

#### 2.2.1 Longitudinal language profiles

Longitudinal language profiles were created by Saloranta and colleagues (2025). The model consisted of seven language variables measured from the participating children at ages 14 months, 30 months, and five years. The language measures included were: (1) at age 14 months, parental questionnaire MacArthur-Bates Communicative Inventories – Words and Gestures (CDI-I) and (2) at age 30 months, Words and Sentences (CDI-T) (Fenson et al., 1991), (3) at age 30 months, the INTERGROWTH-21^st^ Neurodevelopmental Assessment (INTER-NDA) (Fernandes et al., 2020) of which the subsection of language was included, as well as at age five years, (4) Reynell Developmental Language Scales III (Edwards et al., 1997; Kortesmaa et al., 2001) (5) the Similarities subtest of the Wechsler Preschool Primary Scale of Intelligence (Wahlstrom et al., 2018), (6) mean length of utterance calculated from speech sample, and (7) a Nonword Repetition Task.

Latent Profile Analysis in Mplus (Munthén & Munthén, 1998) was used to divide participants into statistically similar clusters of longitudinal development, which resulted in three categories labeled persistent low, stable average and stable high (Saloranta et al., 2025). Most likely class membership statistic was used to determine group membership of each participant. The developmental characteristics of the profile persistent low resembled the growth trajectory of late talking toddlers growing up to meet the criteria for DLD at age 5. The profile of stable average included children performing near age-calibrated mean throughout the years, while the profile of stable high consists of children whose language skills are the strongest in relation to the two other groups throughout the follow-up from 14 months to five years.

#### 2.2.2 MR image acquisition

All visits were conducted by trained research staff. Both parents provided written informed consent for image acquisition, while verbal consent was acquired from the children. A member of the research team met each family before MR image acquisition to practice imaging situation, and to familiarize each family to imaging procedure. On the day of scan, children practiced imaging protocol with a wooden coil before entering the actual MRI scanner.

Children watched a video of their choosing during the scanning. Earplugs and headphones were provided for hearing protection, while foam padding provided comfort and reduced head movement. Participants were instructed to use a signal ball to pause or stop the scanning at any time needed. For an increased sense of security, a member of the research team and parent stayed in the scanner room during the imaging protocol.

A 3T MRI system (MAGNETOM Skyra Fit; Siemens Medical Solutions, Erlangen, Germany) equipped with a 20-channel head/neck coil was used for scanning. The Generalized Autocalibrating Partially Parallel Acquisition (GRAPPA) technique with a parallel acquisition technique factor of 2 was applied to all image acquisitions to speed up image acquisition process. All the scans were completed within one hour. High-resolution 3D T1-weighted images using the magnetization prepared rapid gradient echo (MPRAGE) technique were acquired with the following sequence parameters: repetition time (TR) = 1,900 ms, echo time (TE) = 3.26 ms, inversion time (TI) = 900 ms, flip angle = 9 degrees, voxel size = 1.0 × 1.0 × 1.0 mm^3^, and field of view (FOV) 256 × 256 mm^2^. More in-depth descriptions on MRI data collection details can be read in previous publications (Copeland et al., 2022; Pulli et al., 2022).

#### 2.2.3 Preprocessing of MRI data

Preprocessing of the data was done with the Computational Anatomy Toolbox (CAT12, version r1363; Gaser et al., 2023). A standard preprocessing pipeline, encompassing normalization, segmentation, quality assessment, and smoothing, was applied with the following specifications: the default tissue probability map provided in SPM was used, affine regularization was done with the European brains template, inhomogeneity correction was medium, affine preprocessing was rough, local adaptive segmentation was performed at medium, skull-stripping was done via the Graph cuts approach, voxel size for spatial normalization of the images was set to 1.5 mm and internal resampling for preprocessing was fixed at 1.0 mm. Dartel was used for spatial registration and the MNI152 template (MNI = Montreal Neurological Institute) was used for both Dartel and Shooting templates.

#### 2.2.4 Whole-brain voxel-level morphometry

The Statistical Parametric Mapping (SPM) (Friston, 2007, https://www.fil.ion.ucl.ac.uk/spm/) software is a toolkit for analyzing imaging data. The software models imaging data and creates statistical parametric maps that have a known distribution under the null hypothesis (Penny et al., 2011). Within SPM, Voxel-based morphometry (VBM) is an analysis method that enables voxel-wise comparison of local brain tissue — gray-matter, white matter, and cerebrospinal fluids — concentration across subjects (Ashburner & Friston, 2000, 2001). VBM converts magnetic resonance imaging data from all subjects into a symmetrical brain template through spatial normalization, segments the different brain tissue types (Ashburner & Friston, 2005). The images are then “modulated” using Jacobian determinants to maintain absolute volumetric data that may have changed due to spatial normalization (Mechelli et al., 2005) and then smoothed. Finally, a General Linear Model is used to calculate voxel-wise statistics to identify regions that differ, for example between groups or correlate with variables of interest (Mechelli et al., 2005). The GLM includes multiple-comparison corrections based on Random Field Theory (Mechelli et al., 2005; Worsley et al., 1996).

In this study, the GM density calculated by VBM refers to proportional volumes of the cortex. It is the ratio of GM compared to other tissue types within a specific region. This measure is influenced by various neurobiological processes, including synaptic density, neuronal size, and the degree of myelination (Mechelli et al., 2005).

### 2.3 Statistical analyses

Descriptive statistics were created with IBM SPSS Statistics 30 (2025). Statistical VBM analyses were conducted in MATLAB R2016b (MathWorks, MA) using Statistical Parametric Mapping, version 12 (SPM12) (http://www.fil.ion.ucl.ac.uk/spm/software/spm12/). The three longitudinal groups of language development were compared to each other on whole-brain VBM metrics to discover if between-groups differences existed on cortical GM density. Group comparisons were conducted as general linear models using the ‘paired t-test’ option in SPM. The threshold for forming clusters was at *p* < 0.01 and false discovery rate was used with *p* < 0.05 to account for multiple testing across clusters. Sex and age at the time of brain MRI scan were used as covariates of no interest in the models. Handedness was excluded from analyses due to low number of left-handed participants.

## 3 Results

The study sample consisted of 54.8 % boys, which resembles the structure of the longitudinal language sample (Saloranta et al., 2025). The sample used in this study did not differ from the longitudinal language model in terms of maternal age at birth, duration of pregnancy or maternal socioeconomic status (SES), see Table 1. Descriptive group statistics of the three language profiles of the study sample can be found in Table 2.

**Table 1:** Comparison of core demographic variables between samples of longitudinal language model and the current study.

|  | Longitudinal language model | Sample used in this study | Corrected *p* value of the difference |
| --- | --- | --- | --- |
| N | 1281 | 166 | - |
| Boys, % | 52.46 % | 54.8 % | 0.543 |
| Maternal age at birth of child, years, mean (SD) | 30.85 (4.40) | 30.50 (4.71) | 0.348 |
| Duration of pregnancy, weeks, mean (SD) | 39.79 (1.64) | 39.80 (1.56) | 0.943 |
| Maternal SES, md | 3 | 4 | 0.211 |

*Note.* Group comparisons were calculated using chi-square test of independence for categorical variables and one sample *t* test statistic for continuous variables. SD = standard deviation, md = median, SES = Socio-economic status, variable combining maternal education and income, 1 being lowest level, 4 the highest.

**Table 2:** Descriptive statistics of the three developmental language groups.

|  | Persistent low | Stable average | Stable high | Statistical difference between groups |
| --- | --- | --- | --- | --- |
| N total | 23 | 63 | 80 | $\chi^2(2) = 30.952, p < 0.000^{***}$ |
| N boys (%) | 15 (65.21%) | 38 (60.32%) | 38 (47.5%) | $\chi^2(2) = 3.503, p = 1.00$ |
| Age at time of scan, days, mean (SD) | 1967.57 (37.01) | 1968.13 (44.02) | 1976.14 (50.28) | $(F(2, 163) = 0.644, p = 1.00)$ |
| Ponderal index, mean (SD) | 14.19 (1.37) | 14.12 (1.10) | 14.02 (1.23) | (F(2, 163) = 0.257, $p$ = 1.00) |
| Duration of pregnancy, weeks, mean (SD) | 39.43 (1.71) | 39.54 (1.68) | 40.11 (1.35) | (F(2, 163) = 3.156, $p$ = 0.45) |
| Birth weight, grams, mean (SD) | 3606.14 (344.24) | 3543.13 (507.82) | 3547.70 (484.32) | (F(2, 160) = 0.149, $p$ = 1.00) |
| 5 min Apgar score, md | 9 | 9 | 9 | $\chi$ (2) = 0.095, $p$ = 1.00 |
| Maternal age at birth, years, mean (SD) | 31.00 (4.84) | 30.78 (4.41) | 30.15 (4.92) | (F(2, 163) = 0.458, $p$ = 1.00) |
| Maternal BMI before pregnancy, mean (SD) | 24.88 (5.66) | 24.04 (4.29) | 23.87 (3.66) | (F(2, 159) = 0.500, $p$ = 1.00) |
| Maternal SES, md | 4 | 4 | 3 | $\chi$ (2) = 0.028, $p$ = 1.00 |

*Note.* Group comparisons were calculated using Kruskal-Wallis test for categorical variables, one-way ANOVA for continuous variables and chi square test of proportions for group sizes. Multiple testing corrections were applied to the $p$ values. \*\*\* = statistical significance at level $p < 0.001$ , SD = standard deviation, md = median, ponderal index = weight in kilograms divided by height in meters cubed, SES = Socio-economic status, variable combining maternal education and income, 1 being lowest level, 4 the highest. Handedness was excluded from analysis due to low number of left-handed participants.

Longitudinal profiles of early language were compared against each other on whole-brain voxel-based morphometry to find out if there were language group differences in brain structures. Group comparisons revealed differences in cortical GM density between longitudinal language profiles of persistent low and stable high. The stable high language profile had higher GM density than persistent low in the right superior frontal gyrus (peak p = 0.003, size: 1493 mm^2^, peak coordinates: (16, 10, 50)) (see Figure 1 and Table 3). No differences were found on GM density between language profiles of persistent low and stable average, nor between stable average and stable high language profiles. We found no differences between any language profiles in GM density of gray matter areas associated with ventral and dorsal language pathways.

**Figure 1:**
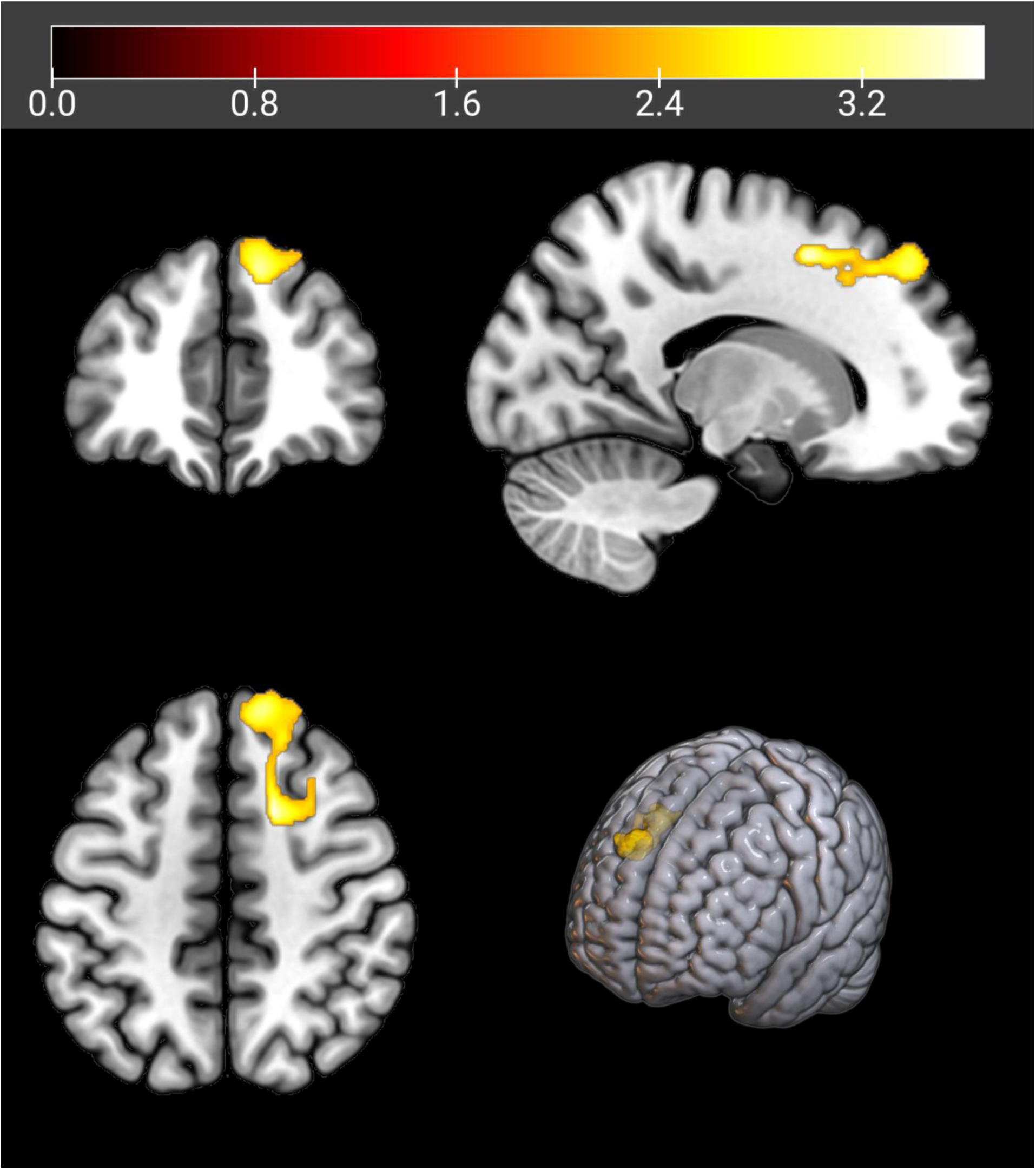
The cluster with higher gray matter density in right superior frontal gyrus for stable high language profile when compared to persistent low. *Note.* The statistics were thresholded at p < 0.01, and FDR corrected at cluster level. The figure was created with MriCroGl software. Image is displayed in neurological convention: right = right side of the brain. The color bar depicts T values.

**Table 3:** Brain regions showing increased density for stable high language profile over persistent low.

| Brain areas (cluster peak location) | MNI coordinates |  |  | Statistic |
| --- | --- | --- | --- | --- |
|  | X | Y | Z | T |
| Right superior frontal gyrus | 16 | 10 | 50 | 3.73 |
| Right superior frontal gyrus | 12 | 44 | 45 | 3.41 |
| Cerebral white matter | 28 | 9 | 42 | 3.11 |

## 4 Discussion

This study aimed to explore differences between longitudinal language profiles of early childhood in GM density at age five in the FinnBrain Birth Cohort. We hypothesized that the differences in early language development would be reflected in differences in cortical areas belonging to the dorsal and ventral language pathways. In contrast to our hypothesis, we found no differences between language profiles and GM density of language pathways. Instead, we found that the children whose language development was strong had denser GM in the right superior frontal gyrus (SFG), in comparison to those whose language development started and stayed on a weak developmental trajectory. We did not find differences in GM density between strong and stable average language profiles, nor stable average and persistent low language profiles. Most notably, no differences were found between any groups in language specific pathways.

The area that we found to differentiate between strong and weak language development is located in “the multiple demand network” (Duncan, 2010). This network of bilateral frontal and parietal cortical areas is responsible for many executive functions, for example inhibition control and attention shifting (for executive functions, see Diamond, 2013; for an overview of neural bases, see Perone et al., 2018). Yeon and colleagues (2024) found these fronto-parietal areas to engage in domain-general processing in their fMRI study, suggesting the network to have an important role in learning new skills. Our finding builds on these theories and findings, suggesting that structural differences in this network are connected to first language acquisition, a complex learning process which requires shifting between cognitive, visual and auditory processes. Children with higher GM density in this network might be better capable of handling cognitive load, which in turn may be reflected better in the language outcomes. This is consistent with a long-standing hypothesis suggesting a close relationship between processing capacity limitations and differences in language development (Federmeier et al., 2020). One way to conceptualize the “capacity limitation” utilizes the concept of executive functions, the set of mental processes that are involved in controlling and regulating behaviors and cognitions (Diamond, 2013). Executive functions are often divided into three main areas: shifting, working memory, and inhibition (Lehto et al., 2003). Shifting refers to the ability to flexibly change between tasks. Working memory stores and manipulates information temporarily. Inhibition refers to choosing reactions and ignoring irrelevant and distracting information while selecting crucial information (Diamond, 2013). Studies suggest that children with DLD exhibit capacity limitations in different components of executive functions (Pauls & Archibald, 2016).

The right SFG that we found to relate to differences in individual differences language development has been found to associate with motor tasks (Martino et al., 2011), working memory execution (Boisgueheneuc et al., 2006; Owen, 2000) and attentional control (Corbetta et al., 2008) in prior studies with functional imaging. More specific results relating to language include association between right SFG and word retrieval (Kemeny et al., 2006), semantic word processing (Gutchess et al., 2010), and verbal short-term memory (Peters et al., 2009). While we acknowledge that neural functions are separate to cortical density in itself, the connection of the right SFG to attention and working memory provides structural evidence for the concept that domain-general learning mechanisms relate to language development trajectories in our data: the children following the strongest language profiles had the largest proportional volumes for the hubs linked to attentional control and working memory. The right SFG is also situated in the Parieto-Frontal Integration Theory neural network proposed by Jung and Haier (2007), a model that presents how interaction between frontal and parietal cortical areas affects differences in individual reasoning. Given that right SFG is responsible for monitoring working memory and word processing, its role in cognitive processes that relate to language is essential. Indeed, verbal working memory is positively associated with language development from ages two to four (Newbury et al., 2016), which could be reflected in structural differences of SFG in our sample.

We argue that our result, together with previous findings of the FinnBrain cohort that link non-verbal cognitive ability to cortical GM volume growth (Pulli et al., 2023), provide a basis for associating cognitive development to early language development through shared principles of neural growth. GM maturation is a heterogeneous process, varying across different cortical areas (Christova & Georgopoulos, 2023; Gogtay et al., 2004). When interpreting the association between gray matter and language, it is important to keep in mind that the direction of the association changes during development. Growth charts of human brain tissue show that global gray matter development metrics (total surface area and total cerebrum volume) peak around ages 11-12 years (Bethlehem et al., 2022), after which they start to decrease, indicating pruning as an effective measure of learning. In children aged five, the growth of gray matter is still in progress, leading to positive correlation between gray matter and language. Language acquisition and abilities have been shown to drive synaptic development and regional neuroplasticity in cortical areas associated with language (Kurth et al., 2018; Sowell et al., 2004). We see that there may be two mechanisms affecting the role of SFG density on language: for one, the formation of new synaptic connections contributing to higher GM density in the SFG, which may result from more developed language skills. Another possible marker of more mature GM tissue could be synaptic pruning. According to Chad and Lebel (2024), adolescent GM volume decrease and density increase associated as synaptic pruning leads to the compactification of glial cells and dendritic branches that survive the pruning process. Although individual synapses are too small for their pruning to directly reduce GM volume, the pruning process may initiate a cascade of changes that ultimately result in significant GM volume loss. Consequently, the lost volume far exceeds the actual volume of the pruned synaptic connections, indicating that pruning contributes to more efficient and compact GM tissue organization (Chad & Lebel, 2024). It might be plausible to argue that a similar process takes place in earlier developmental ages and leads to the higher density observed in the SFG. Overall, our main interpretation is that the higher density in the stable high group can be interpreted as a marker of more mature GM. Given that our language variable extends to years preceding the measurement of GM density, we cautiously propose that gray matter differences present in our data might reflect the extent to which past developmental language trajectory has affected brain structure by age five. However, future studies should continue to test the direction of causality.

One of the main strengths of this study is the use of birth cohort data of general population-based sample before the onset of formal education. In previous studies, participants have been recruited based on their language status, which has resulted in equal sized groups of typical development and language difficulties, be them DLD (Bahar et al., 2024), dyslexia (Bailey et al., 2016) or speech sound disorders (Preston et al., 2014). In contrast, our study used a population-based sample the language status of which was determined by latent clustering after data collection. For this reason, the children exhibiting language difficulties in our cohort are not equivalent to case groups in previous studies. While a strength, we recognize that the data-driven method we used for identifying language difficulties should be interpreted with caution: the model itself includes an amount of classification error.

The children belonging to persistent low language trajectory in our study resemble late talking toddlers that develop DLD later down the line, but the classification is based on statistical modeling of performance on assessments typically used to identify language disorders instead of diagnostic criteria. We chose this approach instead of grouping children based on traditional DLD criteria due to many difficulties in defining this condition discussed by McGregor at al. (2025). We wanted to take advantage of the community-based sample in the FinnBrain cohort study allowing us to include a more representative group of children with observed difficulties in language development across the multiple timepoints and different types of measurements. This comes with some potential statistical limitations. In particular, we relied on most likely class membership for subsequent analyses, despite the low entropy of the language profiles. Because this approach does not account for classification uncertainty, associations with brain measures may have been attenuated and should therefore be interpreted with caution. While acknowledging this limitation, the latent modeling of language development used in our study does extend knowledge of neural correlates of language difficulties to earlier stages of development. Our results are similar to those of Bahar et al. (2024), which suggest that GM structures imageable in late childhood associating to language disorder are observable as early as in age five when basics of language skills have been acquired. As many earlier studies before us (Bailey et al., 2016; Evans et al., 2014; Preston et al., 2014), we found structural differences in areas of right hemisphere to be linked to individual differences in language development, adding to the evidence that central hubs of language processing usually located in the left hemisphere might not be important in emergence of individual differences in early stages of development.

## 5 Conclusion

In this paper, we used a large population-based sample to demonstrate that GM density differs in children belonging to different longitudinal language profiles from one to five years of age. Instead of cortical areas belonging to the ventral and dorsal language streams, differences were identified in the right SFG. Since right SFG is part of the executive functions network, we argue that differences in early language development are associated with this network. Our finding suggests that neural changes linked to language development outcomes are imageable as soon as the basics of language skills have been acquired. Together with the previous finding by Bahar et al. (2024), we suggest that the role of right SFG could prove essential for more detailed differential diagnosis for language disorders.

## Data Availability Statement

Current EU and national legislation on personal data protection of sensitive data, and the informed consents given by the study subjects do not permit open sharing of the data. Investigators interested in research collaboration and obtaining access to the data are encouraged to contact FinnBrain board. Contact information of the Principal Investigators are listed on the project website: https://sites.utu.fi/finnbrain/en/contact/.

## Ethics approval statement

This study was conducted following the guidelines of Helsinki Declaration. Ethics committee statements were acquired from Hospital District of Southwest Finland under the following identification numbers: ETMK 31/180/2011 (neuroimaging), ETMK: 57/180/2011, ETMK: 103/1801/2014, ETMK: 107/180/2012, ETMK 26/1801/2015, ETMK: 59/1801/2013, 26/1801/2015, and ETMK: 63/1801/2017.

## Patient consent statement

The families were informed about the aims of the cohort study as well as about the aims of a specific laboratory measurements before consenting to participate. Parents gave written informed consent for their children’s participation, and usage of register data. Verbal informed consent was acquired from the children prior to MRI scan and language measurements at age five years.

## Acknowledgements

This study would have been impossible to conduct without the participating families. Additional special thanks go to the many researchers, the coordinators and the data team of the FinnBrain Study that enabled the collection and analyses of data. We thank our research nurse Susanne Sinisalo, neuroradiologist Riitta Parkkola, paediatric neurologist Tuire Lähdesmäki and physicist Jani Saunavaara for their role in the 5-year MRI data collection. Special thanks go to Matti Lindberg for creating the socio-economic status variable, and clinical Speech-Language Pathology and Psychology students at the University of Turku for their contributions in collecting the 5-year data on language development.

