## Supplementary Material 1 for "Cortical gray matter density at age five associated with preceding early longitudinal language profiles: A Voxel-based morphometry analysis of the FinnBrain Birth Cohort Study"

### Systematic literature review on whole-brain voxel-based morphometry and early language development

To support the aims of the study, we conducted a systematic literature search on prior studies exploring the connections between brain structures and early language development outcomes by structural MRI, including whole-brain level voxel-based morphometry. This search was then used to conduct a summary of the literature in tabular format.

The systematic review was conducted on PubMed database on the 1st of October 2024 with search phrases "language AND gray matter AND trajectory AND MRI" and "language AND gray matter AND development AND MRI". Studies were filtered by human participants (Child: birth-18 years, Newborn: birth-1-month, Infant: birth-23 months, Infant: 1-23 months, Preschool Child: 2-5 years, Child: 6-12 years, Adolescent: 13-18 years) and written language in English. Inclusion criteria for the studies were 1) focusing on brain structures and early language development and 2) brain structure analyses being conducted in whole-brain level VBM, 3) focus of language is first language acquisition and 4) study is conducted in case-control setting with impaired language development being the case group. Exclusion criteria applied to the articles were 1) region of interest analysis (ROI) that excludes data-driven whole-brain level analyses, 2) participants having developmental impairments other than language disorder, 3) participants having been born preterm, 4) analyses focusing on bilingualism, 5) participants being adults (over 18 yrs). Reading difficulties were accepted as a form of language impairment to extend the pool of potential articles.

Literature search yielded 196 items, 18 of which were duplicates. 178 articles were screened by the title, which resulted in 67 articles being accepted for abstract screening. Of the 67 abstracts, 35 articles were accepted into full-text screening. In total, 11 articles were included in the summary, and the details are summarized in Supplementary Table 1.

**Supplementary Table 1**

*Summary of previous studies exploring the connection between brain structures and early language development by whole-brain level voxel-based morphometry.*

| Study | URL | Group (n) | Mean age in years | Girls (n) | Boys (n) | Scanner field strenght | Analysis software | Threshold for p-values | Correction for p-values | Results overall volume | Results frontal | Results parietal | Results temporal | Results occipital | Results subcortical & cerebellum |
|---|---|---|---|---|---|---|---|---|---|---|---|---|---|---|---|
| Bahar et al. (2024) | https://pubmed.ncbi.nlm.nih.gov/38832358/ | DLD (54) | 12.36 | 15 | 39 | 3.0T | SurfStat | < 0.001 | Monte Carlo simulation of 10,000 iterations and a threshold of p < 0.05 (two-tailed). | Language ability correlated positively with surface area. | DLD < TD: surface area in L & R anterior inferior frontal gyrus (pars triangularis and pars orbitalis). | | DLD < TD: surface area in L & R posterior temporal and ventral occipito-temporal cortex. | | |
| | | TD (74) | 12.52 | 33 | 41 | | | | | No differences in cortical thickness or asymmetry. | DLD < TD: surface area in L & R superior frontal cortex. | | DLD < TD: surface area in L & R anterior cingulate cortex. | | |
| Pigdon et al. (2019)* | https://pubmed.ncbi.nlm.nih.gov/31732792/ | DLD (17) | 10.25 | 7 | 10 | 3.0T | SPM8, VBM8 | < 0.001 uncorrected, < 0.05 FEW-corrected | FWE | | | | | DSD > TD: L inferior occipital gyrus. | DLD > TD: R cerebellum. |
| | | DSD (13) | 10.33 | 10 | 3 | | | | | | | | | | |
| | | TD (40) | 10.25 | 21 | 19 | | | | | | | | | | |

| | | | | | | | | | | | | | | |
|---|---|---|---|---|---|---|---|---|---|---|---|---|---|---|
| Bailey et al. (2017) | [https://pubmed.ncbi.nlm.nih.gov/27324343/](https://pubmed.ncbi.nlm.nih.gov/27324343/) | DYS (14) | 12.5 | 4 | 10 | 3.0T | SPM8 & VBM8 | < 0.05 | Bonferroni | DYS < SRCD & TD: L inferior frontal gyrus. | DYS < SRCD & TD: R posterior cingulate, L supramarginal gyrus. | DYS < SRCD & TD: L inferior and middle temporal gyrus. | DYS < SRCD & TD: R middle occipital gyrus, R lingual gyrus. | DYS < SRCD & TD: L & R thalamus and cerebellum. |
| | | SRCD (11) | 11.5 | 5 | 6 | | | | | DYS > SRCD & TD: R middle frontal gyrus. | DYS > SRCD & TD: R superior parietal lobule, R angular gyrus, R postcentral gyrus. | DYS > SRCD & TD: R inferior temporal gyrus, L superior temporal gyrus. | | DYS > SRCD & TD: L anterior cerebellum, R inferior & anterior & superior cerebellum. |
| | | TD (16) | 11.9 | 7 | 9 | | | | | SRCD > DYS & TD: L middle and inferior frontal gyrus. | SRCD < DYS & TD: R precentral gyrus, R anterior cingulate. | SRCD < DYS & TD: R superior, middle and inferior temporal gyri. | SRCD < DYS & TD: L middle occipital gyrus, R lingual gyrus, R cuneus, L middle occipital gyrus. | SRCD < DYS & TD: R & L inferior cerebellum. |
| | | | | | | | | | | | SRCD < DYS & TD: R postcentral gyrus. | SRCD < DYS & TD: L inferior and superior temporal gyri, | SRCD > DYS & TD: L middle occipital gyrus. | SRCD > DYS & TD: R parahippocampal gyrus. |
| | | | | | | | | | | | SRCD > DYS & TD: R middle cingulate cortex | SRCD > DYS & TD: L superior temporal gyrus, L | | SRCD > DYS & TD: R posterior |

| | | | | | | | | | | | | | | | |
|---|---|---|---|---|---|---|---|---|---|---|---|---|---|---|---|
| | | | | | | | | | | | | and R & L insula. SRCD > DYS & TD: L postcentral gyrus, L supramarginal gyrus, R & L precuneus. | middle and inferior temporal gyrus. SRCD > DYS & TD: R fusiform gyrus, L middle temporal gyrus. | | cerebellum |
| Xia et al. (2016) | https://pubmed.ncbi.nlm.nih.gov/26679527/ | Whole sample | 10-15 | | | 3.0T | SPM8 & VBM8 | < 0.001 uncorrected, < 0.05 corrected | FWE | | DYS < TD in younger and older subgroups: L middle frontal gyrus. | | DYS < TD in younger and older subgroups: L temporo-parietal cortex. | DYS < TD in younger and older subgroups: L superior occipital gyrus. | |
| | | DYS (24) | | 11 | 13 | | | | | | Volume on L dorsal pars opercularis reduced with age for TD but increased for DYS. | | Volume on L ventral occipito-temporal cortex reduced with age for TD but increased for DYS. | | |
| | | TD (24) | | 11 | 13 | | | | | | Younger DYS < younger TD on L dorsal pars opercularis | | | | |

| | | | | | | | | | | | | | | |
|---|---|---|---|---|---|---|---|---|---|---|---|---|---|---|
| | | | | | | | | | | | s, but older DYS > older TD in the same region. | | | |
| | | Younger group | | | | | | | | | | | | |
| | | DYS (12) | 11.0 | 5 | 7 | | | | | | | | | |
| | | TD (12) | 11.0 | 6 | 6 | | | | | | | | | |
| | | Older group | | | | | | | | | | | | |
| | | DYS (12) | 14.1 | 5 | 7 | | | | | | | | | |
| | | TD (12) | 14.1 | 6 | 6 | | | | | | | | | |
| Evans et al. (2014) | https://pubmed.ncbi.nlm.nih.gov/23625146/ | Whole sample | | | | 1.5T & 3.0T | SPM8 & FSL | < 0.001 uncorrected, < 0.05 corrected | FWE | | | TD boys > DYS boys: L supramarginal/angular gyri. TD girls > DYS girls: R precentral gyrus, R central sulcus and L cuneus. | | |
| | | DYS (32) | 9.6 for boys and 10.1 for girls | 17 | 15 | | | | | | | | | |
| | | TD (32) | 8.3 for boys and 9.1 for girls | 17 | 15 | | | | | | | | | |
| Girbau-Massana | https://pubmed.ncbi.nlm.nih | SLI (10) | 9.33 | 4 | 6 | 3.0T | SPM5 | < 0.05 | FDR | TD > SLI | | TD > SLI & SLI+RD: R | | TD > SLI: L & R medial |

| | | | | | | | | | | | | | | | |
|---|---|---|---|---|---|---|---|---|---|---|---|---|---|---|---|
| et al. (2014)* | .gov/24418156/ | | | | | | | | | | | postcentral gyrus. | | occipital gyri. | |
| | | subgroup of SLI, SLI + RD (6) | 9.0 | | | | | | | TD > SLI+RD | | | | SLI > TD: R superior occipital gyrus. | |
| Preston et. al (2014) | https://pubmed.ncbi.nlm.nih.gov/24342151/ | SSE (23) | 9.75 | 5 | 18 | 1.5T | SPM8 & VBM8 | < 0.025 | | | | SSE > TS: L inferior supramarginal gyrus. | SSE > TS: R & L Heschl's gyrus, R & L planum temporale. | TS > SSE: R lingual gyrus. | |
| | | TS (54) | 9.92 | 24 | 30 | | | | | | | | SSE > TS: and R planum polare. | | |
| | | TD (14) | 10.1 | 4 | 10 | | | | | | | | | | |
| Badcock et al. (2012)* | https://pubmed.ncbi.nlm.nih.gov/22137677/ | SLI (10) | 13.5 (8-17) | 1 | 9 | 1.5T | FSL | < 0.001 uncorrected | Not applied | No differences. | SLI > TD: L inferior frontal gyrus. | SLI > TD: L intraparietal sulcus and R insula. | TD > SLI: Posterior superior temporal sulcus bilaterally. | TD > SLI: L occipital pole. | TD > SLI: R caudate nucleus; R substantia nigra. |
| | | TD (16) | 12.5 (6-25) | 9 | 8 | | | | | | TD > SLI: medial frontal polar cortex. | TD > SLI: R medial superior parietal cortex | TD > SLI: R superior temporal gyrus. | | |
| Soriano-Mas et al. (2009)* | https://pubmed.ncbi.nlm.nih.gov/18781595/ | Whole cohort | 5-17 | | | 1.5T | SPM2 | < 0.001 uncorrected, < 0.05 multiple comparisons corrected | Not applied | SLI > TD | | | SLI > TD: R posterior superior temporal gyrus. | SLI > TD: L middle occipital gyrus. | |
| | | SLI (36) | 10.58 | 12 | 24 | | | | | Correlation of age & GM volume in TD but not SLI. | | | | | |

| | | | | | | | | | | | | | | | |
|---|---|---|---|---|---|---|---|---|---|---|---|---|---|---|---|
| | | TD (36) | 10.88 | 12 | 24 | | | | | | | | | | |
| | | Younger cohort | | | | | | | | | | | | | |
| | | SLI (19) | <11 | | | | | | | SLI > TD | | SLI > TD: L motor cortex & precuneus. | SLI > TD: entorhinal area bilaterally; L temporopolar cortex. | | SLI > TD: L caudate nucleus. |
| | | TD (14) | | | | | | | | | | | | | |
| | | Older cohort | | | | | | | | | | | | | |
| | | SLI (17) | >11 | | | | | | | No group differences. | | | | | |
| | | TD (22) | | | | | | | | | | | | | |
| Jäncke et. al (2007)* | [https://pubmed.ncbi.nlm.nih.gov/17010420/](https://pubmed.ncbi.nlm.nih.gov/17010420/) | DLD (21) | 8.33 (4-10) | 7 | 14 | 1.5T | SPM99 | < 0.05 | Not applied | No group differences. | No group differences. | No group differences. | No group differences. | No group differences. | No group differences. |
| | | TD (21) | 9.26 (4-10) | 7 | 14 | | | | | | | | | | |
| Herbert et al. (2003)* | [https://onlinelibrary.wiley.com/doi/epdf/10.1111/1467-7687.00291](https://onlinelibrary.wiley.com/doi/epdf/10.1111/1467-7687.00291) | DLD (24) | 8.3 | 8 | 16 | 1.5T | semi-automated | NA | NA | DLD > TD | | | | | |
| | | TD (30) | 9.1 | 15 | 15 | | | | | | | | | | DLD < TD: Cerebral cortex & caudate nucleus bilaterally smaller in relative size. |

*Note.* * = Information identical to Table 1 in Bahar et al. (2024), DLD = developmental language disorder, DSD = developmental speech disorder, TD = typical development, SLI = specific language impairment, RD = reading difficulty, DYS = dyslexia, SRCD = specific reading comprehension deficit, SSE = speech sound error, TS = typical speech, FWE = family-wise error, FDR = false discovery rate.
